# Digital whole-community phenotyping to assess morphological and physiological features of plant communities in the field

**DOI:** 10.1101/2022.04.27.489638

**Authors:** Verena Zieschank, Robert R. Junker

## Abstract

Traits link observable patterns in plants to ecosystem functions and processes and help to derive general rules and predictions about responses to environmental gradients, global change and perturbations. Ecological field studies often use manual low-throughput methods to assess plant phenotypes and integrate species-specific traits to community-wide indices. In contrast, greenhouse or lab-based studies, mostly in agriculture, employ high-throughput phenotyping for plant individuals to track their growth or fertilizer and water demand. We customized an automated plant phenotyping system (PlantEye F500, Phenospex, Heerlen, The Netherlands) for its mobile application in the field for digital whole-community phenotyping (DWCP). By scanning whole plant communities, we gather, within seconds and non-invasively, multispectral and physiological information while simultaneously capturing the 3-dimensional structure of the vegetation. We demonstrated the potential of DWCP by tracking plant community responses to experimental land-use treatments over two years. DWCP captured short- and long-term changes in morphological and physiological plant community properties in response to mowing and fertilizer treatments and thus reliably informed about changes in land-use. In contrast, manually measured community-weighted mean traits and species composition remained largely unaffected and were not informative about these treatments. Thus, DWCP proved to be an efficient method to measure morphological and physiological characteristics of plant communities, complements other methods in trait-based ecology, provides indicators of ecosystem states, and may help to forecast tipping points in plant communities often associated with irreversible changes in ecosystems.

## Introduction

Trait-based ecology has emerged as a tool to derive general rules and predictions about functional changes in communities in response to environmental gradients, global change components and other perturbations such as species loss or invasions (de Bello et al., 2021; Funk et al., 2017; Lavorel & Garnier, 2002). Quantitative measurements of traits complementing species inventories thus provide the linkage to ecosystem functions and processes (Cadotte et al., 2013; Díaz et al., 2007; Laliberté & Legendre, 2010). For instance, the distribution and variability of traits in grassland communities was related to primary productivity (Liu et al., 2015; Mason & de Bello, 2013; Turnbull et al., 2013), ecosystem stability (Turnbull et al., 2013), and the diversity of organisms in higher trophic levels that interact with the plants (Junker et al., 2015). One major advantage of trait-based assessments of plant diversity is the transferability and thus generalization of findings to other plant communities that may be composed of a different set of species: communities experiencing the same environmental conditions have been shown to converge in species traits despite differences in taxonomic composition (Fukami et al., 2005), indicating that functional traits are of key importance in community ecology and reveal mechanisms complementary to taxonomical information in community assembly (HilleRisLambers et al., 2012; Meiners et al., 2015).

Three major sources of trait data are usually utilized that differ in scale, resolution, and in the way they capture local adaptations and intraspecific variability: 1) Data bases such as the Plant Trait Database TRY (Kattge et al., 2020) that provide species-specific trait values but no information on local adaptations and intraspecific variability. These data bases led to unprecedented insights into the ecology and evolution of plants across taxonomic boundaries (Diaz et al., 2016) and to numerous macroecological studies revealing plant responses to large-scale biotic or abiotic gradients (e.g. Kuppler et al., 2020). 2) Remote sensing by satellites or unmanned aerial vehicles (UAVs) provides large-scale spatial and temporal data, but in a low to medium resolution, respectively (Cavender-Bares et al., 2020; Sun et al., 2021; Wachendorf et al., 2018; Zhu & Woodcock, 2014). Such methods are used for the classification of land-cover types or to predict different aspects of diversity. Additionally, they are used in ecosystem monitoring to create habitat maps or capture characteristics of plant communities (Blackburn et al., 2021; Cruzan et al., 2016). While there are many different types of sensors to be used with satellites and UAVs to record different aspects of plant diversity, cover, and status (Barbedo, 2019), they only provide a two-dimensional representation of the vegetation. 3) Direct field measurements of plant traits using caliper rules, scales, office scanners, and handheld spectrometers may be the most common and direct way of assessing the functional composition and diversity of local plant communities (de Bello et al., 2021; Escudero et al., 2021; Junker et al., 2019; Kuppler et al., 2017; Pérez-Harguindeguy et al., 2016). These measurements are specific to the location and the phenotyped species and individuals and precisely measure selected traits. As these approaches are associated with a high workload, feasible sampling schemes often cannot include all species present in a community and will, to a certain extent, sacrifice some aspects of trait variability (de Bello et al., 2021).

Whereas “low-throughput” phenotyping methods dominate ecological field studies, in agricultural research and applications “high-throughput phenotyping” became standard (Dhondt et al., 2013; Rosenqvist et al., 2019). Adopting such methods for ecological studies may provide novel insights and a more complete view on the phenotypic properties of plant communities and may fill the gap between direct field measurements and airborne remote sensing. We suggest digital automated phenotyping of plant individuals and communities as a novel source of quantitative trait data in ecological field studies that provides complementary data on a different scale while reducing the workload compared to classic trait measurements. A variety of approaches are available to acquire phenotypic raw data such as cameras for RGB color recording, multispectral units for reflectance measuring in the visible, infrared, or near-infrared spectrum, lasers that measure distance for 3D imaging and thermal sensors, among others (Barbedo, 2019; White et al., 2012). The application of such devices under field conditions is often limited because either individual plants need to be transported to a stationary scanner (plant-to-sensor) or scanners are installed in greenhouses or outdoor facilities where they move over a defined set of plants (sensor-to-plant) (Busemeyer et al., 2013; Demidchik et al., 2020; Fiorani & Schurr, 2013). Thus, plant scanners are required that are mobile and therefore applicable under field conditions, independent of a stationary infrastructure.

In this study we used an automated plant phenotyping system (PlantEye F500, Phenospex, Heerlen, The Netherlands) to collect high-resolution multispectral and physiological information on plant communities while simultaneously also capturing the 3-dimensional structure of the vegetation. This method will be referred to as ‘digital whole-community phenotyping’ (DWCP) in the following. Such high-resolution sensors have been predominantly used in the fields of agronomy and ecophysiology to explore the link between genotype and phenotype by scanning plant individuals (Pieruschka & Schurr, 2019).

We customized a MicroScan device (Phenospex, Heerlen, The Netherlands) equipped with the scanner PlantEye F500 in such way that it is fully mobile, can be mounted in the field without limitations, and runs on a mobile battery. This system generates 3D point clouds complemented with multispectral information, and allowed us to track short-term responses of plant communities to experimental land-use changes in a common garden. The common garden contained grass sods originating from three regions of the Biodiversity Exploratories, a long-term research platform in Germany to study the effects of land use on biodiversity and ecosystem processes (Fischer et al., 2010). In the common garden in Marburg, we subjected each of the sods to one of four land-use treatments of differing intensity. We scanned each sod of the common garden multiple times over two growing seasons (*n* = 11 sampling events) recording a time-series of morphological and physiological changes in the plant communities. We extracted a total of 14 parameters from each scan: nine morphological parameters derived from the 3D point cloud and five physiological variables derived from multispectral information. To compare the results of DWCP with classical methods, we additionally manually measured quantitative vegetative traits of plant species on all sods. Using random forest analysis and data from digital and manual phenotyping as well as vegetation analyses, we classified each sod in each sampling event to its provenance and its experimental land-use treatment. Our results demonstrate the potential of automated plant phenotyping systems in assessing morphological and physiological traits and responses to environmental factors of whole plant communities in the field.

## Material & Methods

### Digital whole-community phenotyping

We used the automated plant phenotyping system PlantEye F500 (Phenospex, Heerlen, The Netherlands) to gather multispectral and structural information about plant communities. The PlantEye is equipped with an active sensor that projects a laser line with a wavelength of 940nm vertically onto the vegetation and records the reflection of the laser and the reflectance in Red, Green, Blue, and Near-Infrared with an integrated tilted camera (Kjaer & Ottosen, 2015). All 2D height profiles that are captured by moving the scanner over the plants, driven by an electric motor on a linear spindle axis, are then batched to generate a 3D point cloud. Each data point contains information on the 3D position in a coordinate system (X, Y, Z coordinates) as well as the reflection of red, green, blue, and near-infrared wavelengths (red = 620-645 nm, green = 530-540 nm, blue = 460-485 nm, near-infrared = 820-850 nm). Point clouds are processed with the built-in software HortControl (Phenospex, Heerlen, The Netherlands) that provides morphological and physiological parameters (Table 1). The software also visualizes the 3D point clouds allowing for a quick assessment of scans and the spatial distribution of structural and physiological parameters within the scans (Fig. 1).

**Table 1:** Parameters from digital whole-community phenotyping calculated by the software HortControl (for further information see https://phenospex.helpdocs.com/planteye/planteye-parameters). Definition of parameters are given as well as their functional importance in plant ecology. Information on the interpretation of high and low values of the parameters is given. Finally, the potential range as well as the mean and the 95% confidence interval of our measured data is provided.

| Name | Description | Functional importance | Small values | High values | Potential range | Mean value | 95% CI | References |
| --- | --- | --- | --- | --- | --- | --- | --- | --- |
| **Digital biomass [cm<sup>3</sup>]** | standing biomass; height*3D leaf area | proxy for plant productivity | low biomass | high biomass | 0 - ∞ | 31,459,627 | 30,752,299 - 32,166,955 | Quétier et al. (2007), Guo (2007) |
| **Height [mm]** | average plant height of the community; range from soil to the average top 10% of plants | proxy for plant productivity, nutrient availability, competition for light and growth strategy | lower (maximum) plant height | larger (maximum) plant height | 0 - ∞ | 296.256 | 289.705 - 302.808 | Falster & Westoby (2003), Quétier et al. (2007), Moles et al. (2009) |
| **Height max [mm]** | highest scanned point of the community; range from soil to the highest point of plants |  |  |  | 0 - ∞ | 538.317 | 532.661 - 543.973 |  |
| **Leaf area [cm<sup>2</sup>]** | 3D leaf area; digital leaf area corrected for leaf inclination | proxy for heat load, water retention and gas exchange | small total leaf area | large total leaf area | 0 - ∞ | 107,655.70 | 106,669.4 - 108,642.0 | Stubbs & Wilson (2004) |
| **Leaf area index [cm<sup>2</sup>/cm<sup>2</sup>]** | leaf area per unit sector size; (Leaf Area)/(sector size) | related to photosynthetic activity, respiration and rainfall interception | community of low structural complexity | community of high structural complexity | 0 - ∞ | 0.652 | 0.645 - 0.658 | Fang et al. (2019), Knops & Reinhardt (2000) |
| **Leaf area projected [cm<sup>2</sup>]** | 2D leaf area; amount of sector size covered by leaves | proxy for plant development, growth and competition | sparse plant foliage | large area covered by leaves | 0 - sector size | 53,832.12 | 53,080.02 - 54,584.22 | Lati et al. (2011) |
| **Leaf angle [°]** | average angle of leaves; average angle of the leaf surfaces to the perpendicular | proxy for light capture and water retention, indicator for plants' strategies in dealing with abiotic factors | 0° = vertical leaf | 90° = horizontal leaf | 0 - 90° | 32.928 | 32.703 - 33.153 | Craine et al. (2001), Stubbs & Wilson (2004) |
| **Leaf inclination [cm<sup>2</sup>/cm<sup>2</sup>]** | expresses how erected leaves are in average; (Total Leaf Area)/(Projected Leaf Area) |  | value of 1 = horizontal leaf | higher values = more vertical leaf | 1 - ∞ | 1.880 | 1.868 - 1.891 |  |
| **Light penetration depth [mm]** | deepest point the laser can reach through the canopy; distance between the averaged bottom 20% and top 10% of vertical laser depth | measure for vegetation density and thus proxy for resource availability and above-ground competition | low vegetation density | high vegetation density | 0 - ∞ | 192.251 | 187.447 - 197.055 | Wilson & Tilman (1993), Quétier et al. (2007) |
| **NDVI**<br>**(Normalized**<br>**Digital**<br>**Vegetation**<br>**Index)**<br>**[unit-less]** | ratio of reflected red light to near-infrared light; $(NIR - RED)/(NIR + RED)$ | proxy for vegetation type and<br>productivity, proxy for growth<br>parameters like leaf area index and<br>biomass | -1 – 0 = dead plants or non-plant<br>objects, 0.1 – 0.2 = bare soil, 0.2 –<br>0.5 = unhealthy/sparse<br>vegetation, > 0.5 = healthy/dense<br>vegetation | | -1 - 1 | 0.497 | 0.493 -<br>0.501 | Myneni &<br>Williams (1994),<br>Senay & Elliott<br>(2000),<br>Thenkabail et al.<br>(2000), Kaur et al.<br>(2015) |
| **NPCI**<br>**(Normalized**<br>**Pigments**<br>**Chlorophyll**<br>**ratio Index)**<br>**[unit-less]** | ratio of red and blue channel; $(RED - BLUE)/(RED + BLUE)$ | associated to the chlorophyll content of a<br>plant and thus proxy for the plants'<br>physiological state | lower red<br>reflectance $\triangleq$<br>higher<br>chlorophyll<br>content (0.15-0.25 = green<br>vegetation | higher red<br>reflectance $\triangleq$<br>lower<br>chlorophyll<br>content (>0.25<br>= stressed<br>vegetation) | -1 - 1 | 0.152 | 0.150 -<br>0.155 | Peñuelas et al.<br>(1994), Bannari et<br>al. (2007) |
| **PSRI (Plant**<br>**Senescence**<br>**Reflectance**<br>**Index)**<br>**[unit-less]** | ratio of red & green to near-infrared<br>channel; $(RED - GREEN)/(NIR)$ | associated to the carotenoid/chlorophyll<br>content and thus proxy for leaf<br>senescence | higher<br>chlorophyll<br>content (-0.1 to<br>-0.2 = green<br>vegetation) | lower<br>chlorophyll<br>content (> 0.2<br>= senescent<br>vegetation) | -1 - 1 | 0.133 | 0.130 -<br>0.137 | Merzlyak et al.<br>(1999), Ren et al.<br>(2017) |
| **Greenness/**<br>**Green leaf**<br>**index**<br>**[unit-less]** | shows the relation of the reflectance in<br>the green channel to the red & blue<br>channel; $(2 * G - R - B)/(R + G + B)$ | proxy for combined effects of leaf<br>physiology and canopy structure | more green<br>vegetation | less green<br>vegetation | -1 - 1 | 0.240 | 0.237 -<br>0.243 | Keenan et al.<br>(2014) |
| **Hue**<br>**[°]** | the visible color dependent on the<br>wavelength of light being reflected;<br>expression of color taking into account all<br>three measured colors (RGB) but<br>independently from saturation and<br>brightness value | proxy for the level of photosynthetic<br>pigments and nutrient concentration in<br>plant leaves | green vegetation $\sim 120^\circ$ ($0^\circ$-<br>$60^\circ$=red, $60^\circ$-$120^\circ$=yellow, $120^\circ$-<br>$180^\circ$=green, $180^\circ$-$240^\circ$=cyan,<br>$240^\circ$-$300^\circ$=blue, $300^\circ$-<br>$360^\circ$=magenta) | | $0^\circ$ - $360^\circ$ | 93.653 | 92.987 -<br>94.319 | Adams et al.<br>(1998), Putra &<br>Soni (2017) |

**Figure 1:**
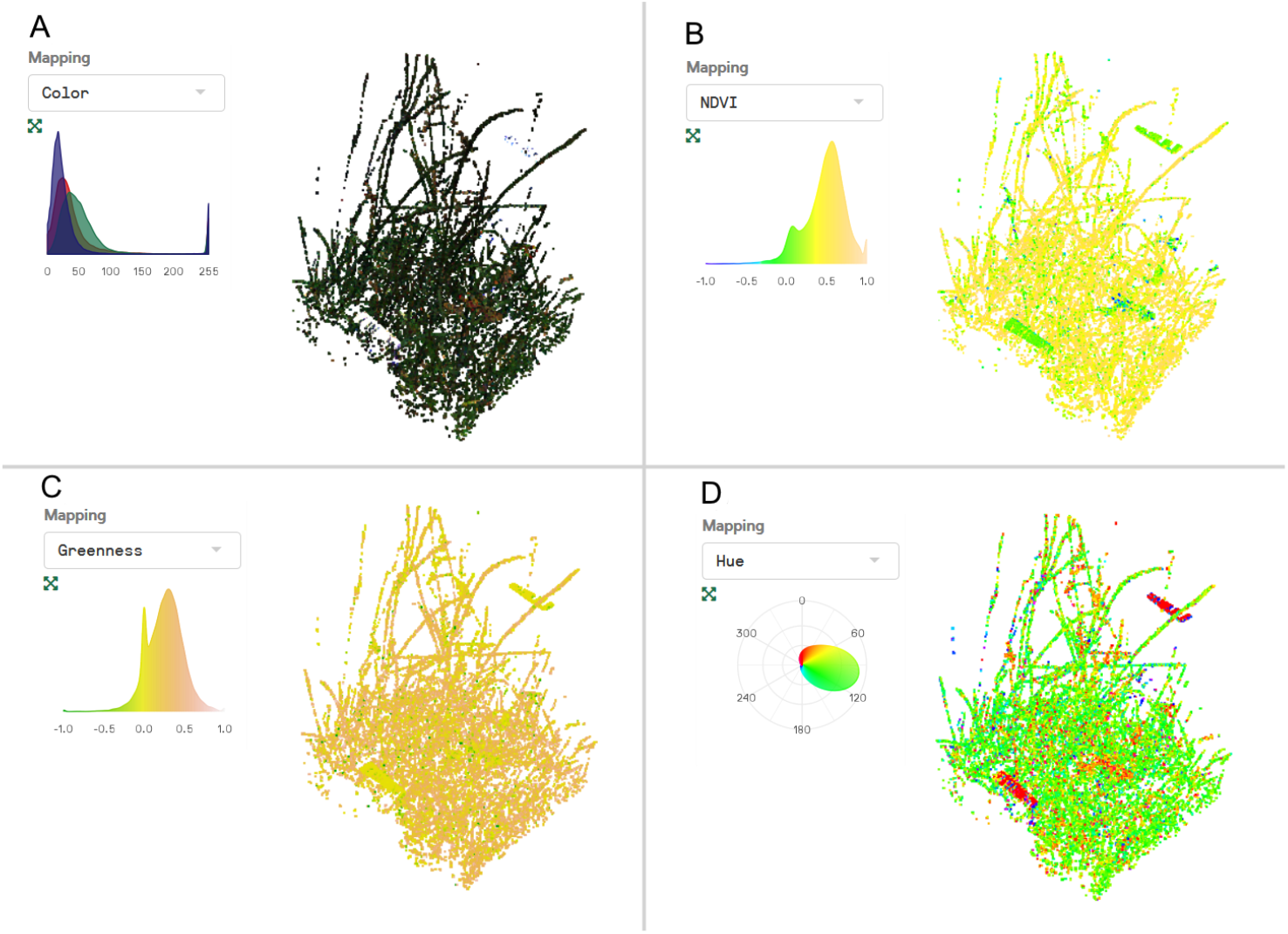
Visualization of the 3D point cloud of a single scan from digital whole-community phenotyping (DWCP) with the built-in software HortControl (Phenospex, Heerlen, The Netherlands). Each point within the cloud contains information on the position (X, Y, Z coordinates) as well as the reflection of red (620-645 nm), green (530-540 nm), blue (460-485 nm) and near-infrared (820-850 nm) wavelengths. Based on this information, RGB color (A), NDVI (B), greenness (C) and hue [°] (D) can be visualized (more options available in HortControl). The distribution of data is shown in the histograms (A-C) or in the color wheel where hues are arranged in a circle (D). The sod originates from plot AEG 09 from the Exploratory Schwäbische Alb and the scan has been made in July 2020 (scan event 1, CW 26).

The PlantEye is independent from light conditions and sufficiently rain-and dustproof for outdoor use. The software processing the scans corrects the intensity of the reflected light for the distance between the vegetation and the sensor using the 3D data. In this study, each scan covered an area of 450 mm width and 300 mm length and 700 mm in height. Each scan had a resolution of <1 mm/pixel, comprising between 155,947 and 425,210 (mean ± sd = 263,692 ± 44,779.07) data points per scan.

For mobile usage, the PlantEye can be mounted onto a mobile frame (MicroScan, Phenospex, Heerlen, The Netherlands) that we customized for use in the field (Fig. 2). We replaced the rear vertical bar with a lightweight carbon tripod (Rollei C6i, RCP Handels-GmbH & Co. KG, Germany), allowing for a much easier positioning on uneven ground while also minimizing the impact on the vegetation. The second vertical bar is replaced by a more robust and lightweight aluminum square tube. Such a tube also stabilizes the horizontal bar that contains the linear spindle axis, providing a better distribution of the weight of the scanner. A solid and waterproof housing contains the electronic control unit, where the horizontal speed of the scanner can be adjusted and scans can be started. An external portable battery was used as outdoor power supply (Polaroid PS600, Polaroid International B.V., 1013 AP Amsterdam, The Netherlands). The final re-built frame had a height of 126,7 cm, a length of 122 cm and a width of 50,7 cm at the stands (Fig 2). A start and stop barcode, 3D printed with customized adjustable brackets to clip them to the square tubes, inidcate the start and the end of the scanning area and also served as reference points to define the above-ground height (Fig. 2).

**Figure 2:**
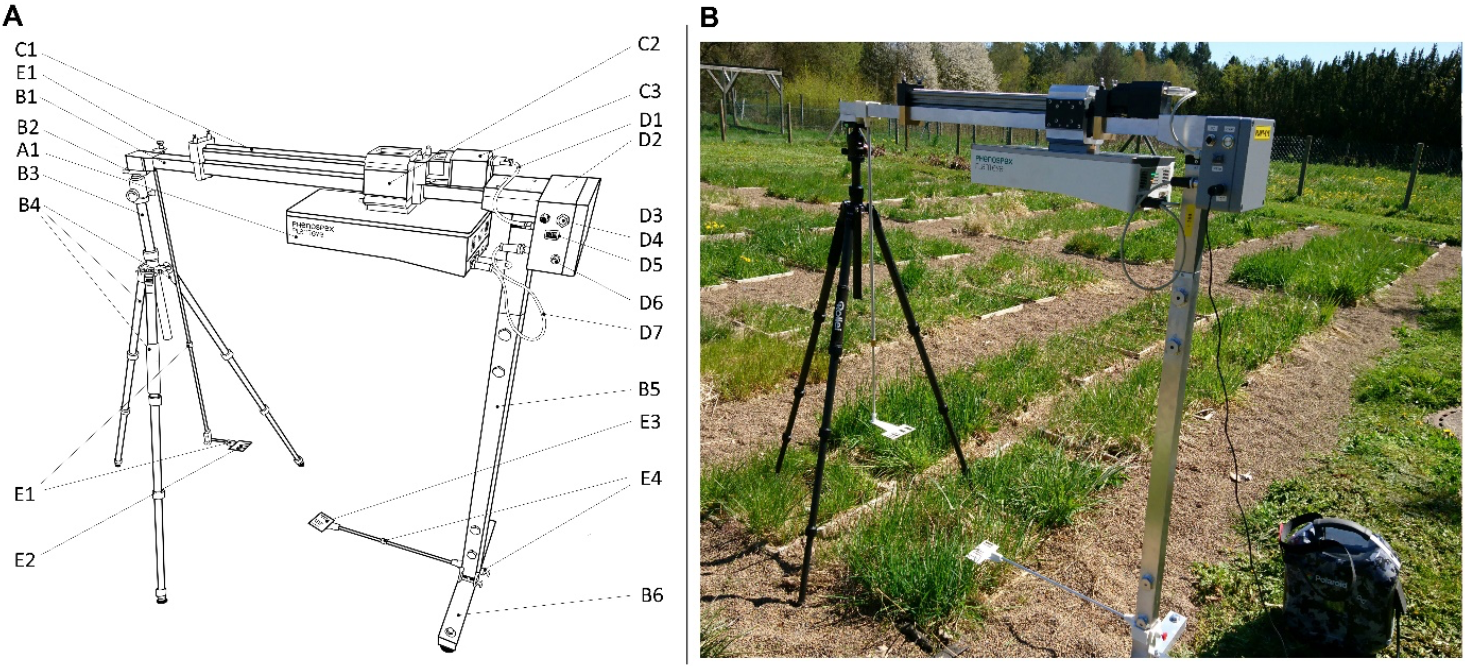
Customized mobile frame allowing the use of the PlantEye plant scanner in the field. **A**) Construction plan of the customized plant scanner: A1) plant scanner, B1) horizontal bar, B2) mounting of the tripod, B3) carbon tripod, B4) length-adjustable legs, B5) vertical bar, B6) rotatable foot, C1) electric linear spindle axis, C2) slide, C3) motor, D1) motor power connection, D2) housing for electronic control unit, D3) port for computer connection, D4) start button, D5) speed control, D6) power cable connection, D7) scanner connection cables, E1) adjustable stop barcode bracket, E2) stop barcode, E3) start barcode, E4) adjustable start barcode bracket. Dimensions of the assembled frame: height = 126,7 cm, length = 122 cm and a width = 50,7 cm. **B**) Photograph of the customized plant scanner for digital whole-community phenotyping in the field.

### Plant communities

To test the potential of digital phenotyping in plant and community ecology, we used plant communities arranged in a common garden established in the Botanical Garden of the University of Marburg, Germany. It was established in April to May 2020 and contains 156 sods. The grass sods originate from the three long-term research sites of the Biodiversity Exploratories (DFG Priority Programme 1374) located in the Biosphere Reserve Schwäbische Alb (ALB) in south-west Germany, the National Park Hainich (HAI) and its surroundings in the center and the Biosphere Reserve Schorfheide-Chorin (SCH) in the northeast of Germany (for further detail about the design see Fischer et al., 2010). We collected sods selected to best represent a land-use gradient from all three regions that were then split into four parts of 50 × 50 cm. Each part was subsequently randomly assigned to one of four experimental land-use treatments. Sods that received the same treatment were arranged in one of three treatment blocks resulting in overall twelve blocks (see supporting information Fig. SI1) (further details about the set up can be found in the supporting information).

Treatments differed in mowing frequency and application of fertilizer in a full factorial design: ‘Control’ (= mowing once per year), ‘Mowing’ (= mowing twice per year), ‘Fertilizer’ (= mowing once per year and fertilizing once per year) and ‘Mowing & Fertilizer’ (= mowing twice per year and fertilizing once per year). Land-use treatments started in mid-July 2020 where all sods were mown and sods assigned to ‘Fertilizer’ and ‘Mowing & Fertilizer’ treatments were fertilized. Mowing was performed using an electric grass cutter (Cordless Grass Shear DUM604RFX, Makita Werkzeug GmbH, Ratingen, Germany), which was used to cut the vegetation to about 2cm in height. The second mowing of sods assigned to ‘Mowing’ and ‘Mowing & Fertilizer’ treatments took place at the beginning of September 2020. In 2021, sods assigned to ‘Mowing’ and ‘Mowing & Fertilizer’ treatments were mown at the end of May and the fertilizing of sods assigned to the respective treatments was done right after. The second mowing in 2021 of all sods in the common garden took place in September. Per sod assigned to a fertilizer treatment, we applied 10.3 g fertilizer granulate (Yara Bela Sulfan, YARA GmbH & Co. KG, Dülmen, Germany), which corresponds to 99 kg N ha^-1^ year^-1^.

### Scanning and manual trait measurements

Over the course of 2020 and 2021, we scanned each plant community in the common garden eleven times (*n* = 11 scan events; 2020: CW 26, 31, 35, 37, 40, 45; 2021: CW 17, 23, 27, 32, 36) using the customized PlantEye scanner. For the scans we used the following software settings in HortControl: pot height = 0, start barcode Z = 170, stop barcode Z = 420, stop barcode y = 300, unit length = 300, length offset = 0, unit width = 450, width offset = −225, height start barcode = 150, height stop barcode = 400. The start barcode was positioned 15 cm above ground, the end barcode was positioned 40 cm above ground. Scanning started five weeks after the common garden was established and was performed in a randomized order that differed in each of the scanning events in order to prevent a spatial and temporal bias in the data. Scan data were complemented with data on the taxonomic composition of the sods as well as with manual measurements of plant traits. Therefore, vegetation surveys were conducted in June/July 2020 and 2021 where all vascular plant species were identified and their cover was estimated with a resolution of 1%. In 2020 we recorded trait values for all plant species that covered >5% in a sod, including plant height [cm], leaf length [cm] and width [cm], leaf dry weight [g] and specific leaf area [mm^2^ mg^-1^] following the protocol of Junker et al. (2020; Junker & Larue-Kontić, 2018). Plant height was measured in the field to the nearest 1 mm using a folding rule. We collected up to five leaves from one to three individuals per species and sod to determine the respective leaf area by scanning the leaf using an office scanner (Perfection 2400 Photo scanner, Seiko Epson Corporation, Nagano, Japan). Leaf area was calculated from the digital leaf scans by dividing the number of pixels per leaf by the number of pixels of a reference square centimeter as obtained from the GNU Image Manipulation Program (GIMP), version 2.10.20 (The GIMP Development Team 2020, retrieved from https://www.gimp.org). After scanning, leaves were dried for at least 2 days at 60°C and weighted on a precision scale (Kern ABS80-4N, KERN & SOHN GmbH, Balingen. Germany). The specific leaf area (SLA) was then calculated by dividing leaf area by leaf dry weight. Additionally, total plant biomass of each grass sod was dried separately after mowing for at least 4 days at 60°C, then subsequently weighted (Sartorius L 610-D, Sartorius AG, Göttingen, Germany) for a direct measure of dry weight biomass [g].

### Statistical analyses

All statistical analyses were conducted in R (R Core Team, 2020. R: A language and environment for statistical computing. R Foundation for Statistical Computing, Vienna, Austria. http://www.R-project.org/). To test the validity of PlantEye parameters, we compared the digital biomass resulting from digital whole-community phenotyping with the biomass weighted after removing it from the sods shortly after the scans using Pearsons’ correlation. To test the explanatory power of DWCP, the plant species composition and the community weighted means (CWM) of manual trait measurements, we ran multiple classifications using *“random forest”*, a machine learning algorithm that assigns samples, in this case the sods in the common garden, to predefined groups: provenance (3 Exploratories), land use treatments (4 experimental land-use treatments), mowing (Y/N) and fertilizer (Y/N) in multiple iterations and estimates the importance of each parameter to achieve the best possible classification (Breiman, 2001). We used three sets of explanatory variables to predict provenance, treatment, mowing and fertilizing: 1) 14 parameters from DWCP with scan events as subsets. 2) plant species list with estimated cover for 2020 and 2021. 3) community weighted mean (CWM) of leaf length, leaf width, leaf dry mass, leaf area and specific leaf area, calculated with the R package *FD* (Laliberté & Legendre, 2010) with species-specific trait values and the species abundances from 2020 and 2021, respectively. All random forest classifications were run with the R package *randomForest* (Liaw & Wiener, 2002) with *mtry* = 4 variables randomly sampled at each split of a decision tree and *ntree* = 10,000 trees to grow for all classifications. We used accuracy values calculated from *randomForest* confusion matrices with the R package *caret* (Kuhn, 2020, version 6.0-86) to quantify classification performance. With the function ‘importance’ of the R package *randomForest*, we extracted variable importance to detect variables that improve classification (Cutler et al., 2007).

## Results & Discussion

In total we analyzed *n* = 1679 digital whole-community phenotyping (DWCP) scans of *n* = 156 sods containing *n* = 167 plant species using the customized PlantEye scanner in summer 2020 and 2021. DWCP turned out to be a valid method to non-invasively assess plant community parameters as indicated by the strong positive correlation between the weighted plant biomass removed from sods and the digital biomass extracted from scans (Pearson’s correlation: *t*_248_= 17.476, *p* < 0.01, *R*^2^ = 0.55, see Fig. SI2), supporting results from plant individuals (Laxman et al., 2018). We used the machine learning algorithm random forest to assign sods to either sod origin (three regions of the Biodiversity Exploratories), experimental land-use treatment (four treatments), mowing treatment (mown once or twice), or fertilizer treatment (with or without fertilizer addition) based on the 14 parameters returned by the software HortControl analyzing the scans. Note that the probability of a correct classification by chance differs between these analyses due to different numbers of categories.

Accuracy of classification of sods to their origin (three regions of the Biodiversity Exploratories) was highest shortly after the establishment of the common garden but strongly decreased in the following weeks and remained low throughout the rest of 2020 and the whole season of 2021 (Fig. 3 A). In contrast, species composition was strongly indicative for sod origin, resulting in a high classification accuracy in both 2020 and 2021 (Fig. 3 B). A classification based on community weighted means (CWMs) of plant species’ traits in the sods was also successful in both years but with a reduced accuracy in 2021 (Fig. 3 B). The three regions of the Biodiversity Exploratories differ in plant species composition (Socher et al., 2013), prevalent soil types, climatic conditions and land-use history but cover a similar range of land-use intensities (Fischer et al., 2010; Gilhaus et al., 2017; Klaus et al., 2013). The fast decrease in accuracy of classification based on DWCP indicates that plant communities show fast responses in morphological and physiological properties to land-use treatments (see below) independent of land-use history, soil type and species composition. In contrast, changes in plant species composition usually emerge only a few years after land-use changes (Komatsu et al., 2019; Read et al., 2018) and thus remained largely stable in our experiment. Accordingly, CWMs of species-specific traits reflect the species composition and thus also slowly responded to land-use changes. The relatively high accuracy in classification by DWCP shortly after the establishment of the common garden may reflect the climatic and edaphic differences in the three regions of origin that may result in differences in vegetation height, digital biomass, hue and NDVI (Fig. 4 A-E). In fact, it has been shown that climate and edaphic conditions affect plant productivity (Burke et al., 1998; Hsu et al., 2012; Lobell et al., 2002). Likewise, remote sensing studies demonstrated changes in NDVI (Paruelo et al., 1997; Tucker & Sellers, 1986) and hue (Dreesen et al., 2013) based on these factors. Additionally, the high accuracy of classification to sod origin based on DWCP may also be explained by the different arrival times of sods at the common garden. Sods from the three regions were transported to the common garden between April and May 2020 and thus the sods from different regions recovered for a different period of time (11 weeks for sods from Schwäbische Alb, 8 weeks for the sods from Hainich, 5 weeks for the sods from Schorfheide-Chorin) from transportation prior to the first scan, which may have resulted in differences in vegetation parameters as well.

**Figure 3:**
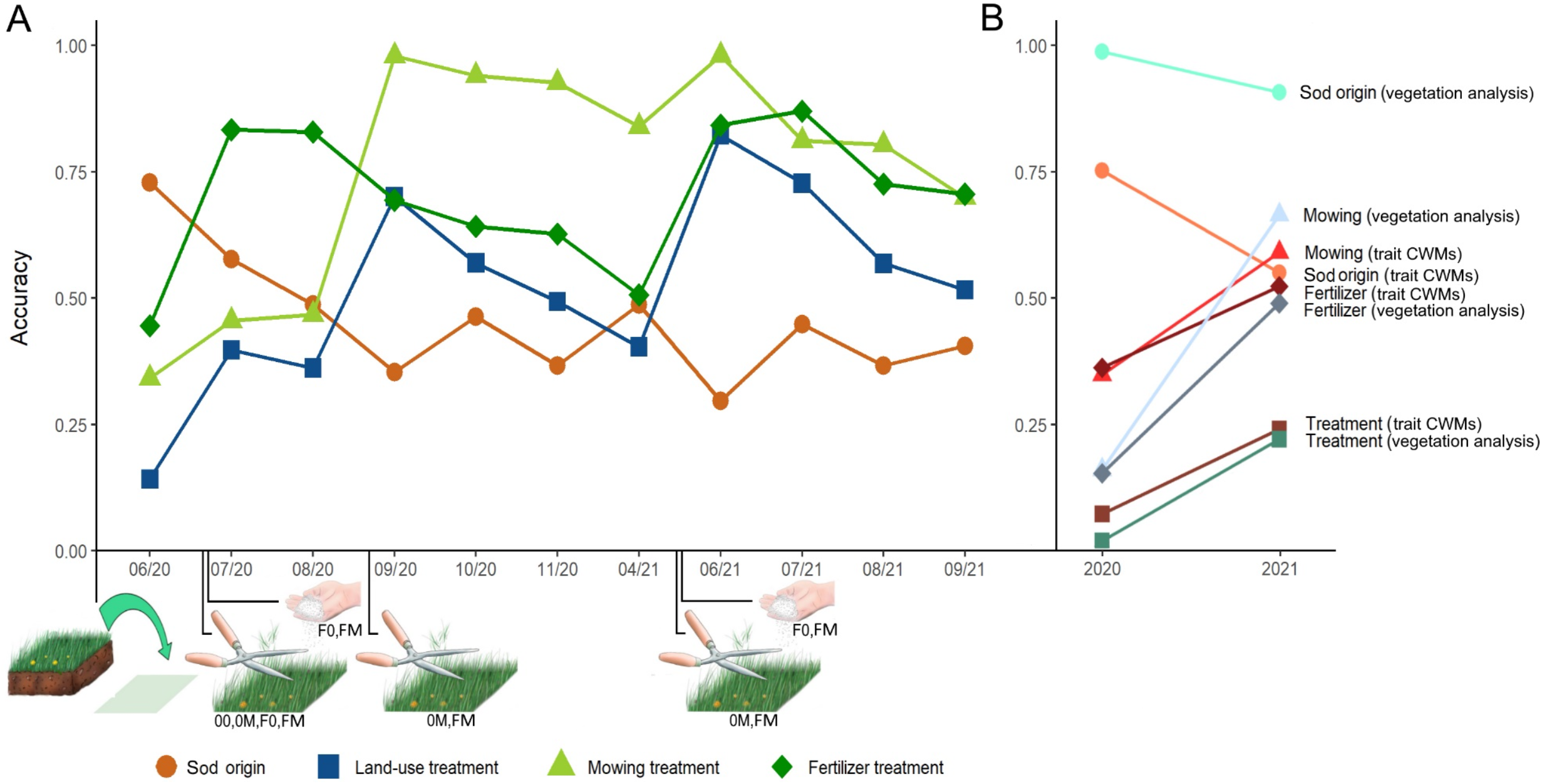
A) Accuracy of random forest classifications (*ntree* = 10,000; *mtry* = 4) of plant communities (= sods) to sod origin (circle, 3 Exploratories), treatments (square, 4 experimental land-use treatments), mowing (triangle, Y/N) and fertilizer (rhombus, Y/N) over the course of 2020 and 2021 with data from digital whole-community phenotyping. Time points of common garden establishment and mowing and fertilizer application events are indicated along the x-axis, treatments affected by these events are indicated below illustrations. **B**) Accuracy of random forest classifications with data from vegetation analysis and community weighted means (CWMs) of traits. As vegetation analyses have been performed only once per year, only changes in accuracy between two years are displayed.

**Figure 4:**
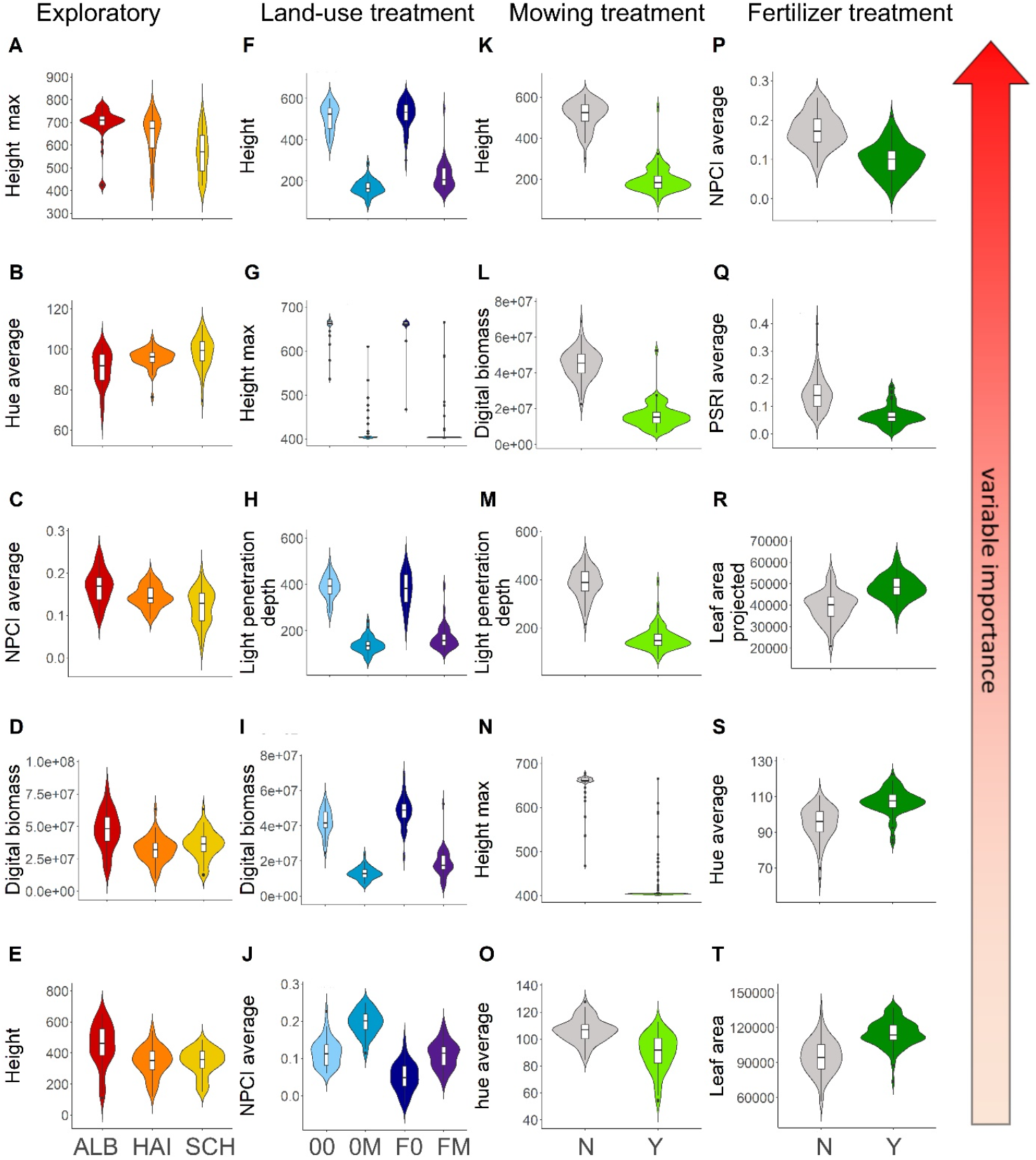
Differences between sod origin (**A**-**E**; 3 Exploratories: ALB = Schwäbische Alb, HAI = Hainich, SCH = Schorfheide-Chorin), land-use treatments (**F**-**J**; 0 = mowing once per year, 0M = mowing twice per year, F0 = mowing once and fertilizing once per year, FM = mowing twice and fertilizing once per year), mowing (**K**-**O**; Y/N) and fertilizer (**P**-**T**; Y/N) in five variables with the highest variable importance for classification in a descending order. For each dependent variable, results of the scan event that received the highest accuracy are shown (scan events 1, 8, 8 and 9, respectively). Boxplots display the distribution of the variables and also show the median, lower and upper quartiles, and outliers. The violin plot outlines depict density distribution of data. All supporting ANOVAs (**A**-**J**) and t-tests (**K**-**T**) were highly significant. Anova and t-test results: **A**) F_2,152_ = 26.16, *p* < 0.01; **B**) F_2,152_ = 16.9, *p* < 0.01; **C**) F_2,152_ = 22.41, *p* < 0.01; **D**) F_2,152_ = 21.25, *p* < 0.01; **E**) F_2,152_ = 16.12, *p* < 0.01; **F**) F_3,148_ = 338.7, *p* < 0.01; **G**) F_3,148_ = 338.4, *p* < 0.01; **H**) F_3,148_ = 228.9, *p* < 0.01; **I**) F_3,148_ = 104.5, *p* < 0.01; **J**) F_3,148_ = 221.3, *p* < 0.01; **K**) *t*_150_= 30.265, *p* < 0.01; **L**) *t*_150_= 23.436, *p* < 0.01; **M**) *t*_150_= 25.659, *p* < 0.01; **N**) *t*_150_= 34.098, *p* < 0.01; **O**) *t*_150_= 8.612, *p* < 0.01; **P**) *t*_152_= 10.813, *p* < 0.01; **Q**) *t*_152_= 9.919, *p* < 0.01; **R**) *t*_152_= −8.866, *p* < 0.01; **S**) *t*_152_= −9.043, *p* < 0.01; **T**) *t*_152_= −9.428, *p* < 0.01.

Accuracies of classifications of sods to the four land-use treatments right after the establishment of the common garden were low for DWCP as well as species composition and CWMs of traits since the land-use treatments had not yet been conducted (Fig. 3 A+B). High peaks in accuracy directly following land-use treatments reveal a clear signal of these treatments detected by DWCP. By the end of the second year the accuracy of classification of sods to the land-use treatments by DWCP parameters has increased considerably, also in scans not directly following the land-use treatments (Fig. 3 A). As stated above, species composition of plant communities can be resistant to land-use change for some years (Komatsu et al., 2019; Read et al., 2018), which is also reflected in the still low, though slightly higher, classification accuracy of treatments from species composition and trait CWMs in 2021. (Fig. 3 B). In contrast to DWCP that allows assessments in high frequencies, vegetation analysis was performed only once per year and trait measurements only once per species. Thus, differences in classification accuracy may also be attributed to differences in the temporal resolution of data collection revealing a clear advantage of DWCP compared to classical methods. In addition, DWCP measures various morphological and physiological parameters, thus providing a multifaceted view on plant communities that are indicative for different land-use treatments (Fig. 4). In remote sensing studies it has been shown that multiple indices outperform single indices in assessing the properties of plant communities (Filella et al., 1995; Hollberg & Schellberg, 2017), which is confirmed by our results. Land-use treatments mostly affected vegetation height, light penetration depth and digital biomass (Fig. 4 F-I), reflecting the strong changes in standing biomass by mowing or fertilizer addition that promotes plant growth and above-ground biomass (Quétier et al., 2007). Note that the parameter ‘maximum height’ is associated with some limitations as it is limited by the maximum height the scanner is able to capture. NPCI as a proxy for the proportion of total photosynthetic pigments to chlorophyll (Bannari et al., 2007; Hollberg & Schellberg, 2017; Peñuelas et al., 1994) was also indicative for land-use treatments (Fig. 4 J). Higher nitrogen availability after fertilizer addition leads to higher chlorophyll concentrations in leaves and consequently to lower NPCI values (Kantety et al., 1996; Schlemmer et al., 2013). In contrast, mowing exposes previously shaded plant parts with lower chlorophyll concentrations (Madison, 1962; Wang et al., 2018) and thus leads to higher NPCI values (Fig. 4 J).

Classification of sods separately to mowing and fertilizer treatments shortly after the establishment of the common garden and prior to the application of the treatments was, as expected, not successful for DWCP, species composition, and CWMs of plant traits (Fig.3 A+B). In both years, the accuracy of classification with DWCP peaked shortly after the respective treatment was conducted, except for the DWCP measurements after the first mowing treatment in 2020 where all sods have been mown. Mowing clearly resulted in structural differences in the plant communities like vegetation height, digital biomass and light penetration depth (Fig. 4 K-N). Strong differences in hue between mown and unmown sods (Fig. 4 O) can be explained by the exposure of previously shaded plant parts with lower chlorophyll concentrations (Guertal & Evans, 2006; Madison, 1962; Wang et al., 2018) and by increased leaf senescence subsequently leading to chlorophyll loss at the cut surface (Howieson, 2001). Therefore, mown sods shifted from green to more yellow colors (Fig. 4 O). Fertilizer treatment was mainly indicated by physiological parameters (Fig. 4 P-T). Fertilizer leads to a higher nitrogen and chlorophyll content in leaves which is measured by both NPCI and PSRI (Bannari et al., 2007; Dong et al., 2012; España-Boquera et al., 2006). NPCI is negatively related to chlorophyll content and thus unfertilized sods had higher values than fertilized sods (Fig 4 P). PSRI increases with degree of senescence due to the resulting loss of chlorophyll and accumulation of carotenoids in leaves (Merzlyak et al., 1999). N fertilizer reduces leaf senescence and promotes higher chlorophyll levels (Dong et al., 2012; Wolfe et al., 1988), both leading to lower PSRI values in fertilized plants (Fig. 4 Q). Hue was slightly higher in fertilized sods (Fig 4 S), indicating overall stronger, more intense green color of plants after the application of fertilizer, most likely initiated by a higher chlorophyll concentration (Adams et al., 1998; Widjaja Putra & Soni, 2017). The higher 2D and 3D leaf area of fertilized sods (Fig. 4 R+T) may be caused by an increase in vegetation density as well as cover density (Quétier et al., 2007) and increased leaf areas (Evans, 1983) after fertilizer treatments. In the first season (2020) after the establishment of the common garden, classification of mowing treatments was more accurate than classification of fertilizer treatments. In the second season (2021), accuracies for both treatments were similarly high (Fig. 3 A), suggesting that the effects of fertilizer treatments that accumulate in plant communities are permanently visible only after a second application. Classification accuracies of sods to mowing or fertilizer treatment based on species composition and CWMs of traits increased slightly from 2020 to 2021 but remained lower than classification by DWCP (Fig. 3 A+B). As mentioned above, plant community composition can be resilient to land-use changes for a period of time (Komatsu et al., 2019; Read et al., 2018) albeit clear differences in species composition due to long-term differences in land-use (Socher et al., 2013). Therefore, short-term responses in vegetation properties as detected by DWCP and measurably in a high temporal resolution may be an indicator for future shifts in species composition and ecosystem properties.

### Future directions

Our data show that digital whole-community phenotyping (DWCP) detects plant community responses to changes in abiotic parameters much earlier than these changes become evident in changes in plant species composition or by using manual trait measurements. Plant traits have been shown to be valuable indicators for ecosystem states (Quétier et al., 2007; Vandewalle et al., 2010) and for tipping points often associated with irreversible changes in ecosystems (Dakos et al., 2019). Likewise, remote sensing data have been utilized to identify grassland land-use intensity focusing on spectral information of pixels (Gómez Giménez et al., 2017). DWCP may bridge the gap between trait measurements in the field and remote sensing and provides a fast and precise method to infer land-use intensity from a combination of morphological and physiological data. Furthermore, the ability to detect community-wide responses to changes in land-use intensity in advance to alterations of species composition and irreversible changes in ecosystems may allow sufficient time to take countermeasures to conserve grasslands.

Future software developments may further increase the potential of DWCP in quantifying community or plant species traits and also as proxies for plant species diversity in communities. The software HortControl, designed to analyze the scans of plant individuals, integrates a whole community and returns community weighted means of the parameters listed in Table 1. Using raw data from digital phenotyping devices, machine learning algorithms succeeded in the segmentation of plant individuals, meaning that morphological structures are recognized in point clouds (Ghahremani et al., 2021; Turgut et al., 2022). In dense grassland plots, this task may be more challenging as individual plants may overlap with other plants and plant parts are not that well separated as in scans of individual plants. Nonetheless, the seven-dimensional raw data (x, y, z position of each point in the point cloud and the red, green, blue, and near-infrared channel) represent a playground for future developments for their analyses and provide the potential for deeper insights into the properties of plant communities.

## Conclusion

Digital whole-community phenotyping turned out to be an efficient method to measure morphological and physiological characteristics of plant communities and thus complements other trait-based approaches and may bridge the gap to remote sensing. Our data suggest that community-wide responses to abiotic parameters are independent on plant species composition and diversity and therefore DWCP represents the next level of generalization attributed to trait-based approaches. Future scans in other ecosystems are, however, required for an assessment of the similarities and differences in plant community responses to land-use changes or other abiotic influences. We conclude that bringing digital plant phenotyping from the lab or greenhouse into the field will reveal detailed insights into the morphological, physiological and functional responses of plant communities in a relevant ecological context.

## Acknowledgements

We thank Gabor Szerencsi and Michael Lietzow for the construction of the customized plant scanner, and Alexander den Ouden and the team at Phenospex for their support. We also thank Annette Schriever, David Meyer, Lilian Winzer, Marie Englert and Marlene Hölzer for help with the fieldwork and data collection and Andreas Titze and the whole team of the botanical garden of the University of Marburg for help with the setup and maintenance of the common garden. We also thank the managers of the three Exploratories, Julia Bass, Anna K. Franke, Miriam Teuscher and all former managers for their work in maintaining the plot and project infrastructure; Christiane Fischer and Victoria Grießmeier for giving support through the central office, Andreas Ostrowski for managing the central data base, and Markus Fischer, Eduard Linsenmair, Dominik Hessenmöller, Daniel Prati, Ingo Schöning, François Buscot, Ernst-Detlef Schulze, Wolfgang W. Weisser and the late Elisabeth Kalko for their role in setting up the Biodiversity Exploratories project. We thank the administration of the Hainich national park, the UNESCO Biosphere Reserve Swabian Alb and the UNESCO Biosphere Reserve Schorfheide-Chorin as well as all land owners for the excellent collaboration. The work has been funded by the DFG Priority Program 1374 “Biodiversity-Exploratories” (Ju 2856/4-1). Field work permits were issued by the responsible state environmental offices of Baden-Württemberg, Thüringen, and Brandenburg.

## Data availability statement

This work is based on data elaborated by the project EXClAvE of the Biodiversity Exploratories program (DFG Priority Program 1374). The datasets (id: 31267 and 31269) are publicly available in the Biodiversity Exploratories Information System (http://doi.org/10.17616/R32P9Q).

## Supporting information

### Common garden “EXClAvE”

The Biodiversity Exploratories are a large-scale research network where the land-use and biodiversity of 50 grassland plots per region have been continuously recorded since 2006 (Fischer et al., 2010). From each of the three regions (“Exploratories”) Schwäbische Alb, Hainich and Schorfheide-Chorin, we selected 13 plots that evenly cover the range of land-use intensity (measured as land-use index LUI, see Blüthgen et al., 2012). In each of the 39 plots, we collected grass sods of the size of 1m^2^ and 10cm thickness in April and May 2020. Every sod was split into four parts of 50×50cm to cover all four experimental land-use treatments and then allocated to one of three treatment blocks that consisted of 14 sods that will be receiving the same experimental land-use treatment: 13 sods from the Exploratories plus one square with steam-sterilized soil to monitor spontaneous plant establishment from the surroundings. Since we have four treatments divided into three blocks each, in total, the common garden consists of twelve treatment blocks (Fig. SI1), adding up to 168 sods.

### Experimental land-use treatments

Sods of fertilizer treatments were treated with 99 kg Nitrogen per ha per year, which corresponds to 10.3 g per sod of the industrial fertilizer ‘YaraBela Sulfan’ (YARA GmbH & Co. KG, Dülmen, Germany), a 24% nitrogen multi-nutrient fertilizer widely used on grassland areas within the Biodiversity Exploratories framework. The fertilizer granulate was evenly distributed on each sod per hand. The amount of fertilizer was calculated from the 90% quantiles of land-use management intensity on the grassland plots of the Biodiversity Exploratories, based on the mean fertilization, mowing and grazing in the years 2006 to 2016, which has also been used to determine the number of cuts per year.

### Preparations prior to the establishment of the common garden and field work specifics

Before the arrival of the grass sods in Marburg, roughly 14 cm of soil were removed from the area of the common garden and subsequently filled again with about 4 cm of gravel to impede upwards root growing. We used wooden frames built from untreated wood with the internal dimension of 50×50 cm as boundaries for the grass sods. After placing the wooden frames, we collected the grass sods from the Schwäbische Alb, National Park Hainich and Schorfheide-Chorin over the course of April 2020 and always planted the sods a maximum of 4 days after collection. In the field, grass sods the size of one square meter were excavated with spades. We then divided them into four equally sized sods (0.25 m2) and transported them to Marburg in plastic boxes. The already harvested sods have been watered during storage until they were planted into the wooden frames in the common garden. During the initial post-planting period sods were sufficiently watered to ensure proper setting of the plants and a good rooting. After the successful planting of all 168 sods, areas between sods were covered with anti-weed fabric before being filled with gravel to the height of the wooden frames. When necessary for survival, grass sods were additionally watered evenly over the whole area of the common garden using two sprinklers (Gardena AquaZoom M, GARDENA GmbH, Ulm, Germany).

## Figures Supporting Information

**Figure SI 1:**
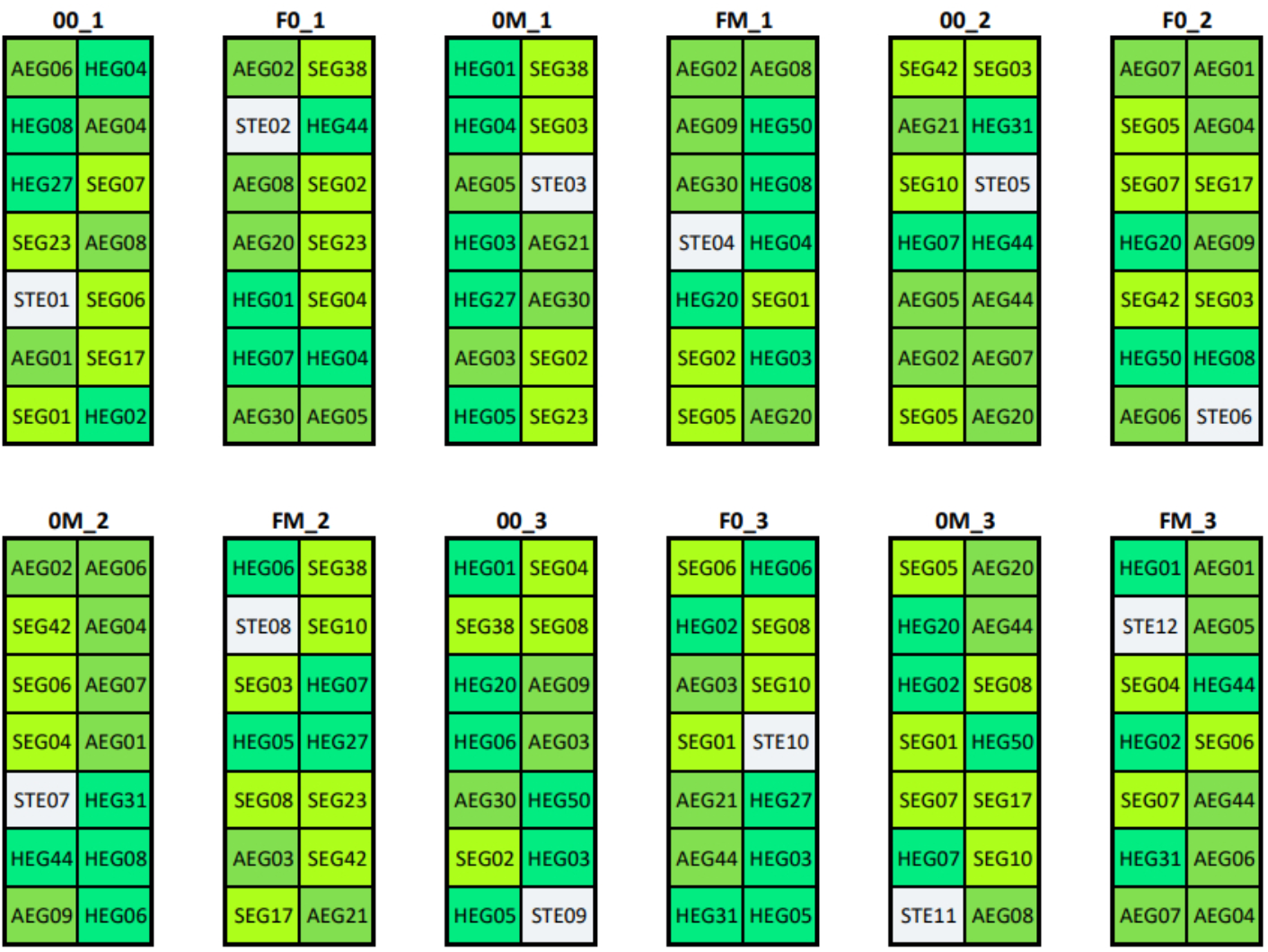
Setup of the common garden at the Botanical Garden of the University of Marburg, Germany. Established April to May 2020, 156 sods originate from the three research sites of the Biodiversity Exploratories (Biosphere Reserve Schwäbische Alb (ALB), National Park Hainich (HAI) and its surroundings, Biosphere Reserve Schorfheide-Chorin (SCH)). Sods were split into four parts of 50 × 50 cm and each part was randomly assigned to one of four experimental land-use treatments (0 = mowing once per year, 0M = mowing twice per year, F0 = mowing once and fertilizing once per year, FM = mowing twice and fertilizing once per year). Sods that received the same treatment were arranged in one of three treatment blocks resulting in overall twelve blocks.

**Figure SI 2:**
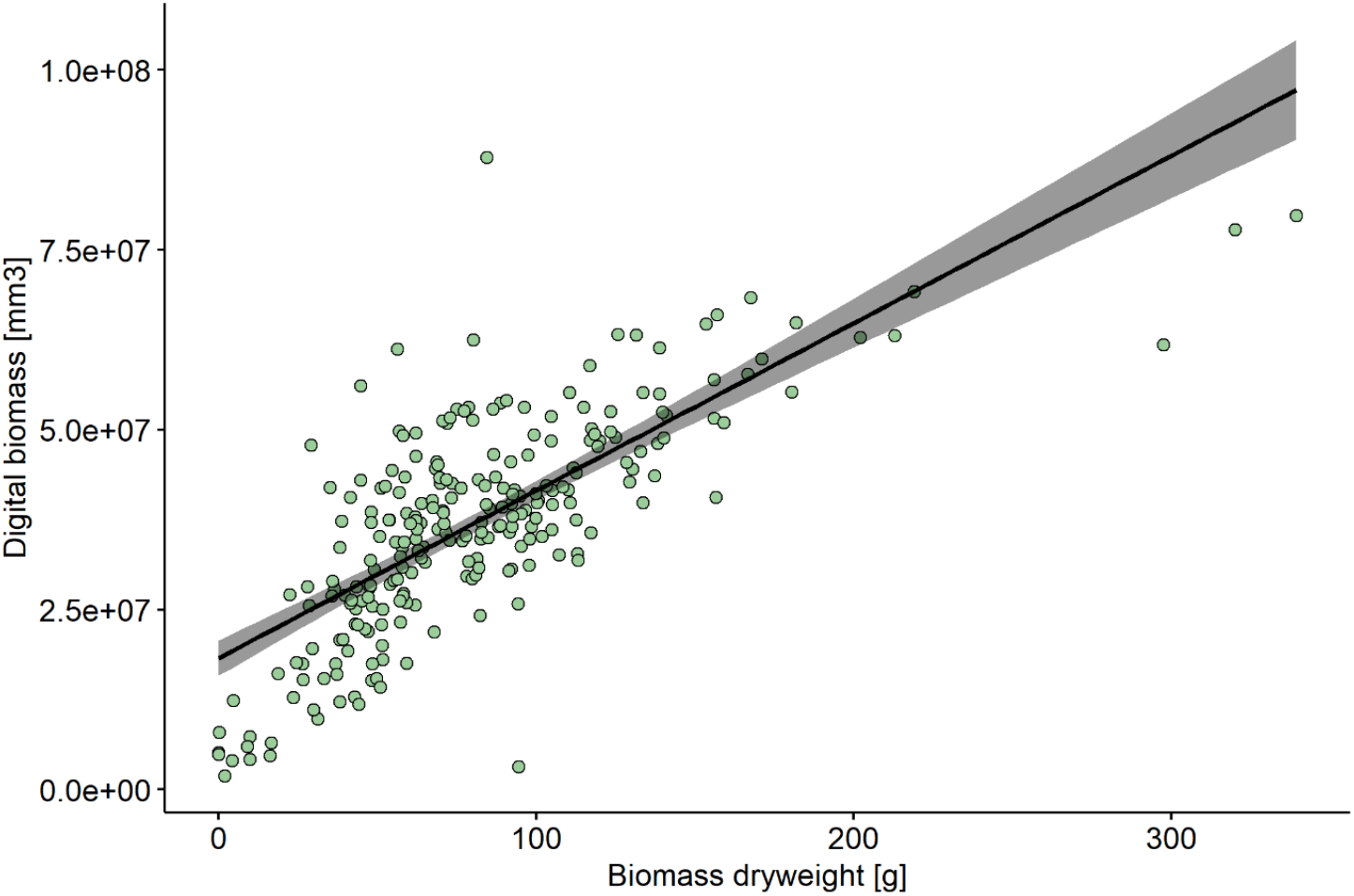
A Pearson product-moment correlation coefficient was computed to assess the relationship between digital biomass as extracted from scans and the weighted plant biomass removed from sods. There was a strong positive correlation between the two variables (*R*^2^ = 0.55, *t*_248_= 17.476) and the relationship was highly significant (*p* < 0.01), confirming that parameters from digital whole-community phenotyping (DWCP) and manual (invasive) measurements are comparable.

